# S100a9 Attenuates Inflammation during Repeated Social Defeat Stress

**DOI:** 10.1101/2022.07.18.500493

**Authors:** Cassandra M. Moshfegh, Safwan K. Elkhatib, Gabrielle F. Watson, John Drake, Zachary N. Taylor, Emily C. Reed, Tatlock H. Lauten, Amelia J. Clopp, Vladimir I. Vladimirov, Adam J. Case

## Abstract

Post-traumatic stress disorder (PTSD), a consequence of psychological trauma, is associated with increased inflammation and an elevated risk of developing comorbid inflammatory diseases. However, the mechanistic link between this mental health disorder and inflammation remains elusive. Using a pre-clinical model of PTSD known as repeated social defeat stress (RSDS), we previously identified that S100a8 and S100a9 mRNA, genes that encode the protein calprotectin, were significantly upregulated in T-lymphocytes after psychological trauma. Calprotectin expression positively correlated with inflammatory gene expression and the mitochondrial redox environment in T-lymphocytes, therefore, we hypothesized that genetic deletion of calprotectin would attenuate the inflammatory and redox phenotype displayed after RSDS. Using pharmacological and genetic manipulation of S100a9 (which functionally eliminates calprotectin) in mice, we unexpectedly observed worsening of behavioral pathology, inflammation, and the mitochondrial redox environment in mice after RSDS compared to wild-type (WT) animals. Furthermore, loss of calprotectin significantly enhanced the metabolic demand on T-lymphocytes suggesting this protein may play an undescribed role in mitochondrial regulation. This was further supported by single-cell RNA sequencing analysis demonstrating that RSDS and loss of S100a9 primarily altered genes associated with mitochondrial function and oxidative phosphorylation. Taken together, these data demonstrate the loss of calprotectin potentiates the RSDS-induced phenotype, which suggests its observed upregulation after psychological trauma may provide previously unexplored protective functions.

## Introduction

Post-traumatic stress disorder (PTSD), an illness characterized by behavioral pathology such as withdrawal, learned helplessness, hyperarousal, and flashbacks, affects nearly 45 million Americans (1-5). Individuals diagnosed with PTSD show significantly elevated risks for the development of comorbid inflammatory pathologies such as autoimmune, metabolic, and cardiovascular diseases (6-16). While inflammation is often observed in individuals with PTSD (13-18), and this may underlie this inherent risk for comorbid inflammatory diseases after PTSD, the mechanistic link between psychological trauma and altered immune function remains unknown and understudied.

Previous work from our laboratory identified a significant elevation in two inflammatory calcium binding proteins, S100a8 (calgranulin A) and S100a9 (calgranulin B), in T-lymphocytes in a mouse model of psychological trauma (*i*.*e*., repeated social defeat stress [RSDS]) (19). Together, these proteins form the heterodimeric pro-inflammatory protein calprotectin, which has been extensively studied within the innate immune system, but had not been reported in T-lymphocytes until our previous work (19). Canonically, calprotectin acts as a damage associated molecular pattern (DAMP), while also sequestering ions such as iron, manganese, zinc, and calcium to inhibit pathogen growth (20-23). Interestingly, extracellular calprotectin has been shown to enhance the development of autoreactive CD8^+^ T-lymphocytes and enhanced IL-17A production in T-lymphocytes (24), and has been implicated in numerous autoimmune diseases (22, 25-28). Furthermore, calprotectin is known to be redox-regulated and plays a critical role in intracellular redox signaling (29). Given that we previously demonstrated that calprotectin correlated with behavioral pathology, inflammation, and redox changes after RSDS (19), these data are strongly suggestive that calprotectin may play a mechanistic role in potentiating the pro-inflammatory T-lymphocyte phenotype we and others have observed after RSDS (19, 30-32).

In the present study, we hypothesized that loss of calprotectin would attenuate the pathology associated with RSDS. To test this hypothesis, we investigated the behavioral, inflammatory, redox, metabolic, and gene expression changes of RSDS in S100a9 knock-out (S100a9^-/-^) mice, which lack functional calprotectin. Surprisingly and in contrast to our hypothesis, we observed that loss of S100a9 exacerbated circulating and T-lymphocyte inflammation, and worsened behavioral pathology after RSDS. Moreover, the mitochondrial redox and metabolic environments of S100a9^-/-^ T-lymphocytes were significantly perturbed compared to WT RSDS mice. Single-cell RNA sequencing analysis on S100a9^-/-^ T-lymphocytes displayed that loss of S100a9 conveyed significant impacts to genes regulating mitochondrial function and oxidative phosphorylation, suggesting a significant mitochondrial role of S100a9 in T-lymphocytes. Together, these data show for the first time a functional role of S100a9 in T-lymphocytes that may be protective in attenuating the global phenotype after psychological trauma.

## Materials and Methods

### Mice

Wild-type (WT) C57BL/6J mice were obtained from Jackson Laboratories (#000664, Bar Harbor, ME, USA). S100a9^-/-^ mice were obtained from the Mutant Mouse Resource and Research Centers at the University of California Davis (#049540-UCD, Davis, CA, USA). CD1 male retired breeder mice were purchased from Charles River at 4-6 months old (#022, Wilmington, MA, USA). All experimental mice (not CD1) were backcrossed to the C57BL/6J background a minimum of 10 generations to create congenic strains, bred in house to eliminate shipping stress and microbiome shifts, and co-housed with their littermates (≤5 mice per cage) prior to the start of experimentation to eliminate social isolation stress. All mice were housed with standard corncob bedding, paper nesting substrate, and given access to standard chow (Teklad Laboratory Diet #7012, Harlan Laboratories, Madison, WI, USA) and water *ad libitum*. Male mice between the ages of 8-12 weeks were used for all experiments, except for juvenile experiments which utilized male and female mice of 3-4 weeks of age. Experimental mice were randomized, and when possible, experimenters were blinded to the respective cohorts until the completion of the study. Mice were sacrificed by pentobarbital overdose (150mg/kg, Fatal Plus, Vortech Pharmaceuticals, Dearborn, MI, USA) administered intraperitoneally. All mice were sacrificed between 7:00 and 9:00 Central Time to eliminate circadian rhythm effects on T-lymphocyte function. All procedures were reviewed and approved by the University of Nebraska Medical Center and Texas A&M University Institutional Animal Care and Use Committees.

### Repeated Social Defeat Stress (RSDS) Paradigm

RSDS utilizes aggressive male retired breeder CD1 mice to exert psychological and physical trauma on experimental male mice of a C57BL/6J background, and was performed as previously described (19). Briefly, CD1 mice were pre-screened for aggression (19). Aggressive CD1 mice were placed into individual clean cages three days prior to starting the protocol. Experimental mice were placed in the home cage of a CD1 mouse to induce a physical confrontation and fear induction for 5 minutes in the absence of food, water, and nestlets. Following this interaction, a clear perforated divider was placed into the CD1’s cage, and both mice remained co-housed for 24 hours. This process was repeated with a new CD1 aggressive mouse daily for 10 consecutive days. Control mice were pair housed in the absence of a CD1 for the duration of the protocol. On day 11, mice were behavioral assessed (described below), and euthanized the following day (day 12) for tissue collection. Mice were continuously monitored for visual wounding and lameness; None of the mice utilized herein met the threshold for exclusion (wounds >1cm or any lameness).

### Social Interaction Test

The social interaction test was performed as previously described (19). The social interaction test consists of an open field chamber (40×40cm, Noldus Information Technology, Leesburg, VA, USA) with a wire mesh cage (6.5×10cm) at one end of the open field. First, both control and experimental mice were individually tested in the enclosure with an empty mesh cage for 2.5 minutes. Then both the control and experimental mice were tested in the enclosure with a novel aggressive CD1 mouse present in the mesh cage for 2.5 minutes. Ethanol and water were used to clean the test chamber between each session. Sessions were recorded and digitally analyzed using Noldus Ethovision XT 13 software (Leesburg, VA, USA). The social interaction zone ratio was calculated by dividing the time the mouse spent in the interaction zone close to the mesh cage in the presence versus the absence of a novel CD1 mouse.

### Elevated Zero Maze

The elevated zero maze was performed as previously described (19). The elevated zero maze test consists of an elevated circular maze (50cm diameter, 5cm track width, Noldus Information Technology, Leesburg, VA, USA), divided into four even quadrants, with two closed arm portions (20cm wall height) and two open arm portions. Both control and experimental mice were individually run on the maze for a period of 5 minutes. Total distance moved along the maze and time spent in the open arms of the maze were assessed using recorded and digitally analyzed footage via Noldus Ethovision XT 13 software (Leesburg, VA, USA).

### Treatment Regimens

Paquinimod (MedChemExpress, #HY-100442, Monmouth Junction, NJ, USA), a S100a9 pharmaceutical inhibitor, was reconstituted in 10% DMSO, 40% PEG300, 50% saline solution and delivered by osmotic minipump (Thermo Fisher Scientific, #NC0551767, Waltham, MA, USA) at a flow rate of 1.5 mg/kg/day. This low dosage was chosen as to minimize off target effects as previously reported (33). Minipumps were surgically implanted subcutaneously on the flank of mice three days before the start of the RSDS protocol. Controls were implanted with vehicle filled osmotic minipumps.

### T-Lymphocyte Isolation and Activation

T-lymphocytes from spleens were isolated using negative selection, as previously described (34). Briefly, splenocytes in a single cell suspension were negatively selected for Pan, CD4^+^, and CD8^+^ T-lymphocytes using the EasySep Mouse T-Cell Isolation Kit, EasySep™ Mouse CD4^+^ T-cell Isolation Kit, and CD8^+^ T-cell Isolation Kit (StemCell Technologies #19851, #19852, #19853, Vancouver, BC, USA), respectively. Cell viability was measured via a Bio-Rad TC20 Automated Cell Counter using trypan blue exclusion, and cell purity was assessed via flow cytometry (average purity achieved is >90%). T-lymphocytes were cultured in RPMI media, with 10% Fetal Bovine Serum, 10mM HEPES, 2mM Glutamax, 100 U/mL penicillin/streptomycin, and 50µM of 2-mercaptoethanol. Cells were activated using Dynabeads Mouse T-Activator CD3/CD28 (Gibco #11452D) at a ratio of 1:1 cells to beads in culture for 72 hours prior to assessment.

### Calprotectin protein assessment

Extracellular calprotectin was measured in plasma and culture media using the Mouse S100a8/S100a9 Heterodimer DuoSet Solid Phase Sandwich ELISA Kit (R&D Systems #DY859605, Minneapolis, MN, USA) as per manufacturer’s instructions.

For intracellular calprotectin assessment, protein was extracted from pan T-lymphocytes using Pierce™ RIPA Buffer (Thermo scientific 89900) supplemented with 1% Halt™ Protease Inhibitor Cocktail (Thermo scientific 1862209). Protein concentration was quantified using Coomassie Plus 200 Bradford Assay Reagent (Fisher PI23238). S100a8 and S100a9 were detected using the Jess Automated Western Blot (Bio-techne). Briefly, samples were diluted to recommended working concentration of 0.4 mg/mL with 0.1x Sample Buffer and 5x Fluoro Mix. Samples were denatured and loaded onto the Jess plate with appropriate reagents and primary antibodies: Anti-S100a8 Polyclonal Antibody – Rabbit (Fisher PIPA579948) diluted 1:20 and anti-S100a9 Polyclonal Antibody – Rabbit (Invitrogen PA5-79950) diluted 1:50. Analysis was performed using Compass Software for Simple Western.

### RNA Extraction, cDNA Production, and Quantitative Real-Time RT-PCR

Select mRNA transcript levels were assessed as previously described (35). Briefly, RNA was extracted from isolated T-lymphocytes using the RNAeasy mini kit (Qiagen #74104, Valencia, CA, USA) according to the manufacturer’s protocol. The concentration of RNA per sample was determined spectrophotometrically using a Nanodrop 2000 Spectrophotometer (Thermo Fisher Scientific, Waltham, MA, USA). One microgram of RNA was then converted to cDNA utilizing the High-Capacity cDNA Reverse Transcription Kit with RNase Inhibitor (Applied Biosystems #4374966, Grand Island, NY, USA). cDNA was then subjected to SYBR green for IL-2, IL-6, IL-17A, and TNFα (Applied Biosystems #4385612, Grand Island, NY, USA) (**Supplementary Table 1**) and FAM PrimerPCR™ Probe Assay mixes for IL-22, S100a8, and S100a9 (BioRad, 12001950, Berkeley, CA, USA) quantitative real-time PCR. The PCR product specificity was assessed by thermal dissociation for SYBR primers, and efficiency assessed for linearity of amplification. A threshold in the linear range of PCR amplification was selected and the cycle threshold (Ct) was determined. Levels of transcripts were then normalized to 18S or RPS18 mRNA loading controls (ΔCT) before determining a relative fold change (2^-ΔΔCT method).

### Flow Cytometry

Flow cytometry was performed as previously described (34). Briefly, spleens were harvested from WT and S100a9^-/-^ control and RSDS mice. For immunophenotyping combined with mitochondrial redox environment assessment, splenocytes were isolated and stained with 1 µM MitoSOX Red (superoxide sensitive mitochondrial-localized probe, Thermo Fisher Scientific #M36008, Waltham, MA, USA) and cell type discriminating fluorescent antibodies CD19 APC-Cy7, CD3e PE-Cy7, CD11c APC, CD11b SB-436, and NK1.1 SB-600 (BioLegend, 115529, San Diego, CA, USA; Thermo Fisher Scientific, 25-0031-81, 17-0114-81, 62-0112-82, 63-5941-82, Waltham, MA, USA) for 30min at 37°C prior to data acquisition. For T-lymphocyte mitochondrial mass assessment, T-lymphocytes were stained with 1 µM MitoTracker Green FM (Thermo Fisher Scientific, M7514, Waltham, MA, USA) and CD4 PE-Cy7 and CD8 BV-786 (BD Biosciences, BDB561479 and BDB563332, Mississauga, ON, Canada) antibodies for 30min at 37°C prior to data acquisition. Cells were analyzed on either a BD LSRII or a Thermo Fisher Attune NxT flow cytometer, and data was analyzed using FlowJo software.

### Plasma cytokine Analysis

Cytokines in plasma were measured as previously described (19). Briefly, cytokine concentrations were assessed using Mesoscale Discovery 29 U-Plex Mouse Biomarker Group (#K15083K-1, Rockville, MD, USA) per manufacturer’s instructions. Analysis was performed using a Mesoscale QuickPlex SQ 120 and analyzed using Mesoscale Discovery software.

### Bioenergetics Analysis

T-lymphocyte mitochondrial bioenergetics were performed as previously described (36). Briefly, splenic T-lymphocytes were plated in Seahorse Bioscience base media (Agilent #102353, Boston, MA, USA) supplemented with 11 mM D-glucose, 1 mM sodium pyruvate, and 1x GlutaMAX. Cells were adhered to Seahorse microplates using 1 μg/cm^2^ Cell-Tak (Corning #354240, Corning, NY, USA). To measure mitochondrial respiration, mitochondrial inhibitors (Seahorse Bioscience Cell Mito Stress Test Kit #103015-100, Santa Clara, CA, USA) were utilized at 1 μM oligomycin, 1 μM 7 carbonyl cyanide-4 phenylhydrazone (FCCP), and 10 μM rotenone/antimycin A.

Glycolytic bioenergetics were performed as previously described (36). Briefly, splenic T-lymphocytes were plated in Seahorse Bioscience base media supplemented with only 1 mM sodium pyruvate, and 1x GlutaMAX. To measure glycolysis in these cells, glycolytic agents (Seahorse Bioscience Cell Glycolysis Stress Test, Agilent #103017-100, Santa Clara, CA, USA) were utilized at 10 mM D-glucose, 1 μM oligomycin, and 50 mM 2-deoxyglucose (2-DG).

### Incucyte Growth Curves

Splenic T-lymphocytes were counted and plated at an optimal seeding density of 8×10^4^ cells per 100 µL in 96 well Microtitration ImageLock plates (VWR, #4379, Radnor, PA, USA). The cells were cultured with anti-CD3 and anti-CD28 activation beads and imaged over the course of 96hrs in the Essen Bioscience IncuCyte S3 Live-Cell imager and analysis system.

### Single-Cell RNA Sequencing

Single-Cell RNA sequencing was conducted as previously described (19). Briefly, three control wildtype, three RSDS wildtype, three control S100a9^-/-^, and three RSDS S100a9^-/-^ male mice were sacrificed, and their spleens were collected for splenocyte preparation. Approximately 2,500 single cells per sample were captured, lysed, and RNA was reverse transcribed and barcoded using a 10x Genomics Chromium instrument and Chromium Single Cell 3’ Reagent Kits v2 reagents (10x Genomics, Pleasonton, CA, USA). Sequence libraries were constructed and then quantified by qpcr using the KAPA Library Quant Kit (Illumina) from KAPA Biosystems (Roche, Pleasonton, CA, USA). All sequence preparation and acquisition was performed at The University of Nebraska Medical Center Genomics Core Facility.

After validating the quality of the single-cell data with FastQC v0.11.9 (37), bioinformatics was conducted on an AWS R4 instance type utilizing Ubuntu 18.04. Alignment was performed by STAR 2.7.10a (38) utilizing GRCm39 primary genome assembly to generate the count data, which was imported into Seurat 4.0 (39) running on R v4.1.2. 113,281 cells remained after quality control metrics (number of genes greater than 200 and less than 2500, mitochondrial content less than 10%). Seurat default parameters were used to log normalize, scale, and integrate all samples together (variable features = 2000, reduction method for integration = reciprocal PCA). In total, Seurat produced 16 unidentified clusters. By first utilizing marker genes and secondly by correlating the first principal component of the RNA assays between clusters identified by Seurat and clusters identified by the authors of the *Tabula Muris* spleen dataset (40), 7 cell types were identified (B, CD8a, CD4, Monocyte/Macrophage, Dendritic, Gamma Delta T, and Natural Killer). Differential expression between phenotypes of a particular cell type employed the Wilcoxon Rank Sum test and used default parameters except for the log fold threshold, which was set to 0.50. In order to detect enrichment against known biological processes, Ingenuity Pathway Analysis (Version 73620684) used with a pre-ranked list of the differentially expressed genes and their corresponding log-fold change and p-values.

### Statistics

A total of 174 animals (87 control, 87 RSDS) were utilized in these studies. While all animals underwent behavioral testing, not all animals could be utilized for each biological or physiological parameter. However, all graphs denote N values for a particular experiment along with each graph displays individual data points. All data sets were assessed for normality using the Shapiro-Wilk test before utilizing a specific statistical test. For graphs with only two cohorts, either a Student’s t-test or Mann-Whitney U-test was performed. For graphs with four cohorts, a 2-way ANOVA was performed, with Tukey’s multiple comparisons test for individual variances computed for each comparison. Simple linear regression analysis was utilized for correlation analyses. Significance was determined with a difference at p<0.05 and exact significant p-values are displayed on individual graphs (non-significant p-values not shown).

## Results

### S100a8 and S100a9 mRNA and Protein are Elevated in T-Lymphocytes after RSDS

Previous work from our laboratory demonstrated that calprotectin was elevated over 3-fold in circulation of RSDS mice, and we were the first to report its presence in T-lymphocytes utilizing single-cell RNA sequencing (19). Herein, we again validate that socially-defeated adult male mice consistently show elevations in S100a8 and S100a9 mRNA transcript levels in splenic T-lymphocytes (**Figure 1A**). Extending these findings, we confirmed that adult RSDS mice possess intracellular S100a8 and S100a9 protein in T-lymphocytes, and these proteins significantly correlate with each other within these cells (r=0.9827, p<0.0001; **Figure 1B**). While knock-out of S100a9 caused the expected loss of S100a9 within T-lymphocytes, interestingly, it also led to a complete loss of intracellular S100a8 (**Figure 1B**). While S100a8 has been shown to have functions independent of S100a9 (and vice versa), in T-lymphocytes it appears they are dependent upon one another for stabilization and function. Furthermore, we found that T-lymphocytes excrete calprotectin, RSDS potentiates production from these cells, and S100a9^-/-^ mice do not produce any detectable secreted calprotectin (**Figure 1C**). To extend this observation to include both sexes, we utilized a juvenile model of RSDS where pre-pubescent mice are exposed to the 10-day RSDS protocol with an aggressive adult male CD1, and then singly housed for four weeks prior to behavioral and physiological analysis. In this model, both male and female mice displayed significantly elevated S100a8 and S100a9 mRNA in T-lymphocytes one month after RSDS (**Supplemental Figure 1**). Taken together, these data report for the first time that T-lymphocytes generate S100a8 and S100a9 protein, and expression of calprotectin is elevated both acutely and chronically following RSDS.

**Figure 1.**
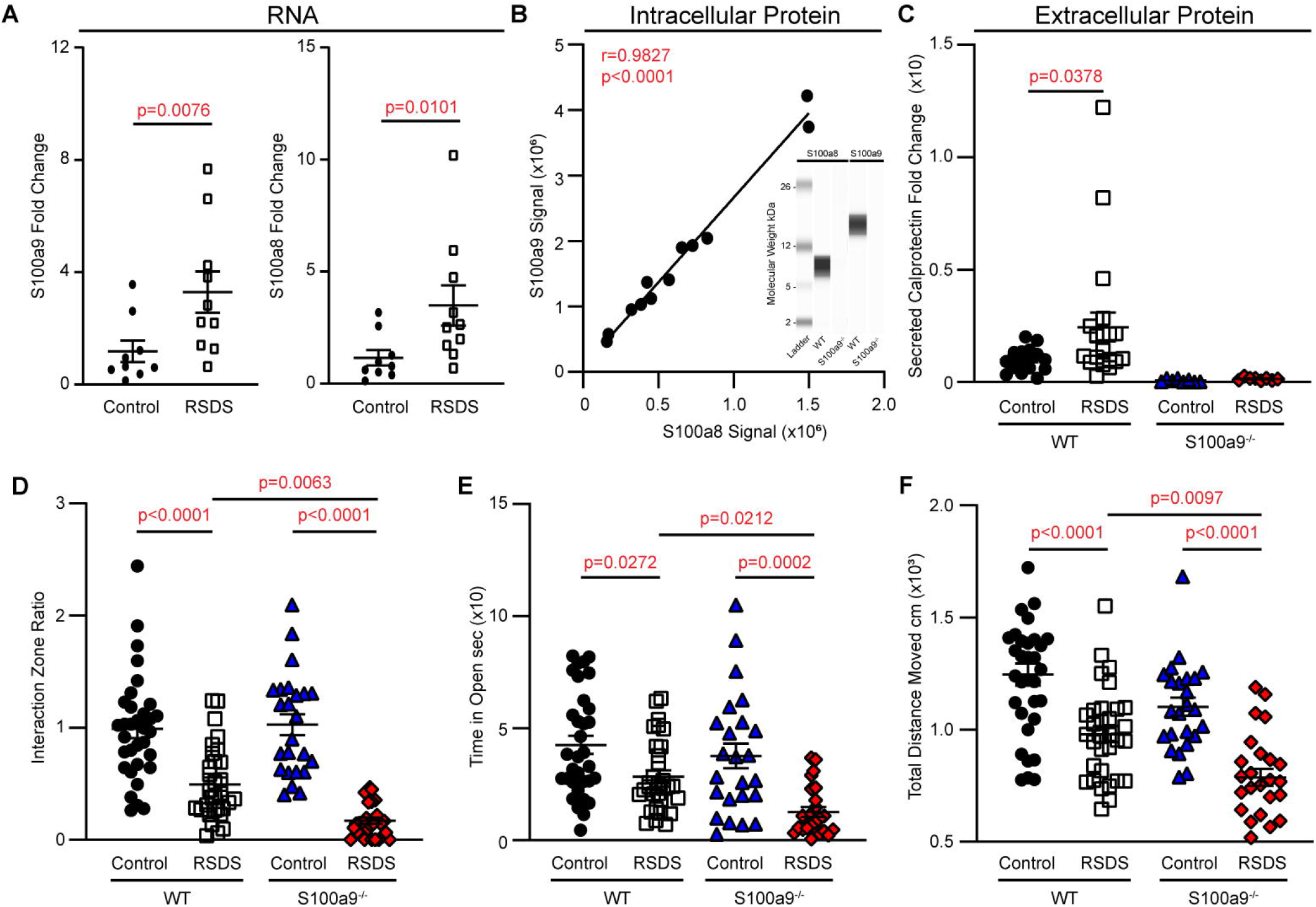
Loss of S100a9 potentiates RSDS-induced behavioral changes. WT and S100a9^-/-^ mice were run through RSDS and behavior testing, after which splenic T-lymphocytes were isolated from these animals. **A**. S100a9 and S100a8 mRNA levels assessed by real-time quantitative PCR in freshly isolated pan T-lymphocytes (n = 9 control, 10 RSDS). **B**. S100a8 and S100a9 intracellular protein quantification by Jess automated western blot analysis on naÏve splenic T-lymphocytes (n = 6 control, 6 RSDS). *Inset*, generated image of S100a8 and S100a9 intracellular protein in a WT and S100a9^-/-^ T-lymphocytes. **C**. Extracellular calprotectin in media of cultured T-lymphocytes assessed by ELISA (n = 6 WT control, 9 WT RSDS, 8 S100a9^-/-^ control, 8 S100a9^-/-^ RSDS). **D**. Quantification of the interaction ratio from social interaction testing. **E-F**. Quantification of the time spent in open arms and total distanced moved from elevated zero maze testing (D-F: n = 33 WT control, 32 WT RSDS, 24 S100a9^-/-^ control, 25 S100a9^-/-^ RSDS). Statistics by Mann-Whitney U-test, Pearson correlation, or two-way ANOVA with Tukey’s post hoc analysis where appropriate.

### Deletion and inhibition of S100a9 Significantly Worsens the RSDS Behavioral Phenotype

To understand if elevated calprotectin played a functional role after psychological trauma, we utilized the S100a9^-/-^ mouse to examine the consequences of calprotectin loss. As expected, RSDS mice showed decreased pro-social behavior with increased anxiety-like behavior in WT animals (**Figure 1D-F**). However, loss of S100a9 showed a worsened phenotype with less variability in these behavioral parameters compared to WT mice (**Figure 1D-F**). Interestingly, all S100a9^-/-^ mice possessed an interaction ratio <1 after RSDS (**Figure 1D**), demonstrating 100% of these animals appear “susceptible” as defined by Nestler and Russo et al. (41, 42). To understand if elevated circulating calprotectin played a functional role after psychological trauma, we pharmacologically inhibited calprotectin by the use of paquinimod. Paquinimod is a clinically-used inhibitor that prevents S100a9 from binding to the receptor for advanced glycation end products (RAGE) and Toll-like receptor 4. Paquinimod infusion significantly worsened the social interaction ratio after RSDS compared with vehicle infused animals, but did not have a significant impact on anxiety-like behavior between the two groups (**Supplemental Figure 2A-C**). Together, these data suggest that loss or inhibition of S100a9 potentiates RSDS-induced behavioral alterations.

**Figure 2.**
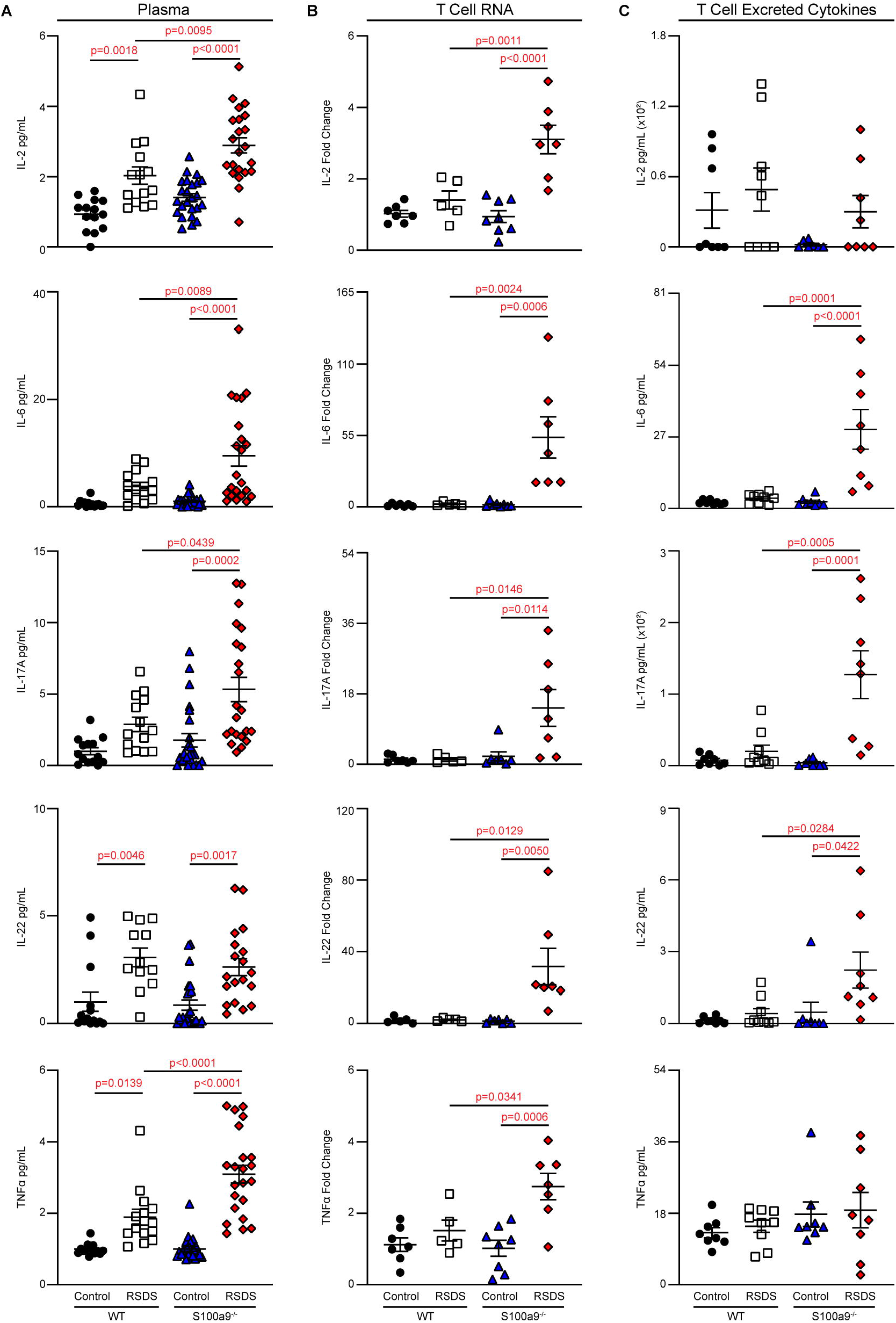
Loss of S100a9 exacerbates the inflammatory phenotype displayed after RSDS. WT and S100a9^-/-^ mice were run through RSDS, after which plasma and splenic T-lymphocytes were isolated from these animals. **A**. Circulating cytokines in plasma assessed by Mesoscale Discovery (n= 14 WT controls, 14 WT RSDS, 24 S100a9^-/-^ control, 22 S100a9^-/-^ RSDS). **B**. T-lymphocyte inflammatory cytokine mRNA assessed by quantitative real-time RT-PCR (n= 7 WT controls, 5 WT RSDS, 8 S100a9^-/-^ control, 7 S100a9^-/-^ RSDS). **C**. Excreted inflammatory cytokines in media of cultured T-lymphocytes assessed by Mesoscale Discovery (n= 8 WT controls, 9 WT RSDS, 8 S100a9^-/-^ control, 8 S100a9^-/-^ RSDS). Statistics by two-way ANOVA with Tukey’s post hoc analysis throughout.

### Loss of S100a9 Exacerbates RSDS-induced Inflammation

We have previously identified a specific subset of circulating inflammatory proteins (*i*.*e*., IL-2, IL-6, IL-17A, IL-22, and TNFα) are induced after RSDS (19, 30-32). Given that S100a9 has numerous reported pro-inflammatory properties, we originally speculated that this inflammatory phenotype of RSDS would be attenuated in S100a9^-/-^ mice. Counter to this hypothesis, we observed that S100a9^-/-^ RSDS mice showed the same or a significantly exacerbated inflammatory profile as compared to WT RSDS mice, whether it be in circulation, freshly isolated T-lymphocyte mRNA levels, or cytokines produced from T-lymphocytes artificially activated *ex vivo* for 72 hours (**Figure 2A-C**). Moreover, exogenous supplementation of calprotectin on unstressed cultured T-lymphocytes attenuated both IL-6 and IL-17A mRNA expression compared to untreated controls (**Supplemental Figure 3A-B**). In summary, we found the loss of S100a9 intensifies inflammation after RSDS, and may in fact play a protective or anti-inflammatory role in T-lymphocytes.

**Figure 3.**
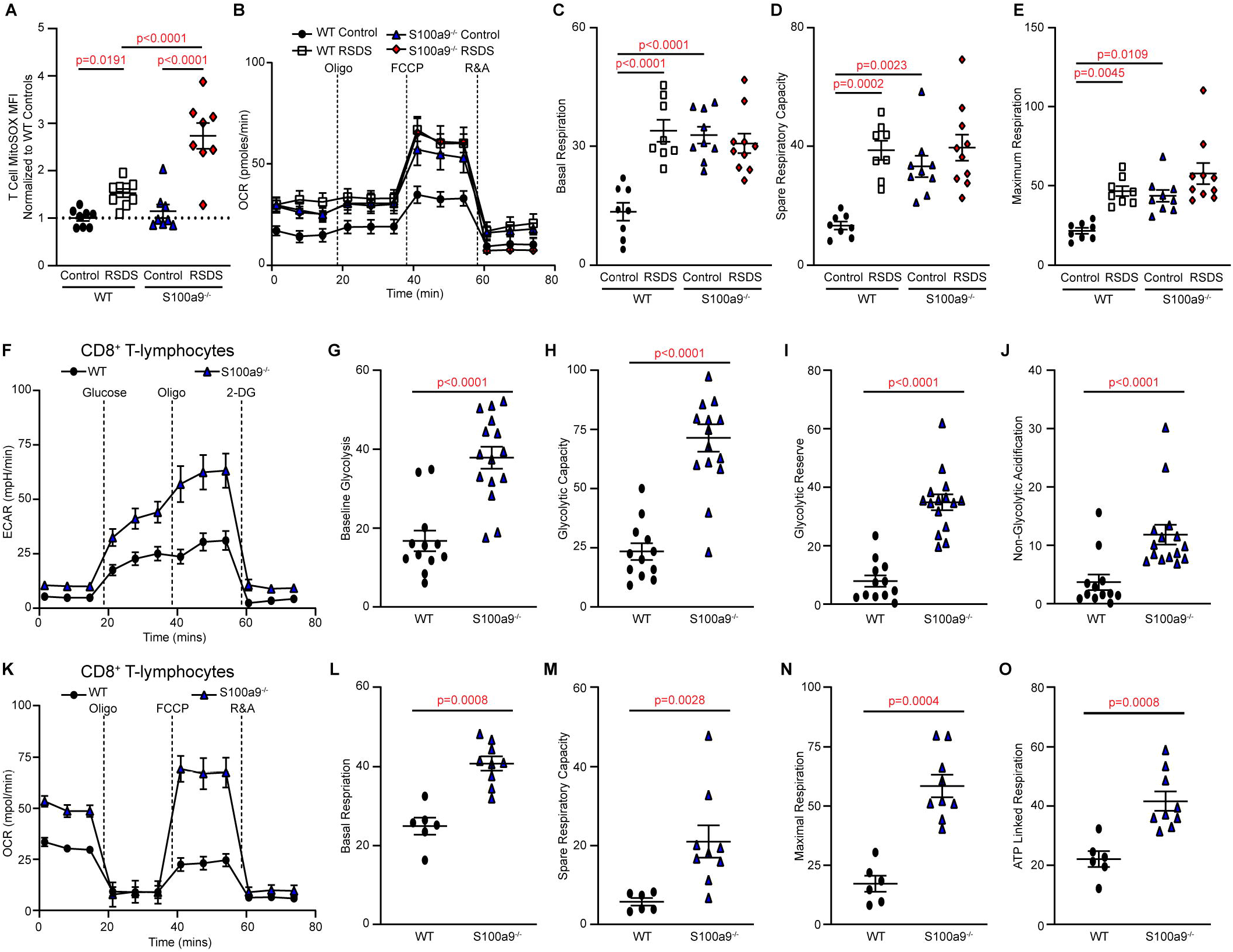
Loss of S100a9 disrupts T-lymphocyte mitochondrial and metabolic homeostasis. WT and S100a9^-/-^ mice were run through RSDS, after which splenic T-lymphocytes were isolated from these animals. **A**. MitoSOX Red mean fluorescent intensity (MFI) assessed by flow cytometry. Data normalized WT control T-lymphocytes to control for interexperimental variance (n= 8 WT controls, 8 WT RSDS, 8 S100a9^-/-^ controls, 8 S100a9^-/-^ RSDS). **B**. Representative oxygen consumption rate (OCR) curve of mitochondrial stress test. (n= 6 WT control, 5 WT RSDS, 9 S100a9^-/-^ control, 10 S100a9^-/-^ RSDS). **C-E**. Quantification of various metabolic states from the mitochondrial stress test. **F**. Representative extracellular acidification rate (ECAR) curve of a glycolysis stress test (n=12 WT, 15 S100a9^-/-^). **G-J**. Quantification of metabolic states from the glycolytic stress test. **K**. Representative oxygen consumption rate (OCR) curve of mitochondrial stress test (n= 6 WT, 9 S100a9^-/-^). **L-O**. Quantification of metabolic states from the mitochondrial stress test. Statistics by Mann-Whitney U-test or two-way ANOVA with Tukey’s post hoc analysis where appropriate.

### Loss of S100a9 Alters Mitochondrial Redox and Metabolism in T-lymphocytes

Our lab has previously shown that mitochondrial superoxide and metabolism regulates T-lymphocyte inflammation, and that T-lymphocyte mitochondrial superoxide is elevated following psychological trauma (19, 34, 36). We again confirmed this phenomenon herein in WT animals, but found that the loss of S100a9 further potentiated mitochondrial superoxide in T-lymphocytes after RSDS (**Figure 3A**). Paquinimod infusion mirrored this potentiation of T-lymphocyte mitochondrial superoxide (**Supplemental Figure 4A**). Moreover, exogenous supplementation of calprotectin in culture decreased mitochondrial superoxide levels in both CD4^+^ and CD8^+^ T-lymphocytes compared to untreated controls (**Supplementary Figure 4B**). Together, these data suggest calprotectin plays a previously unidentified role in regulating the mitochondrial redox environment in T-lymphocytes.

**Figure 4.**
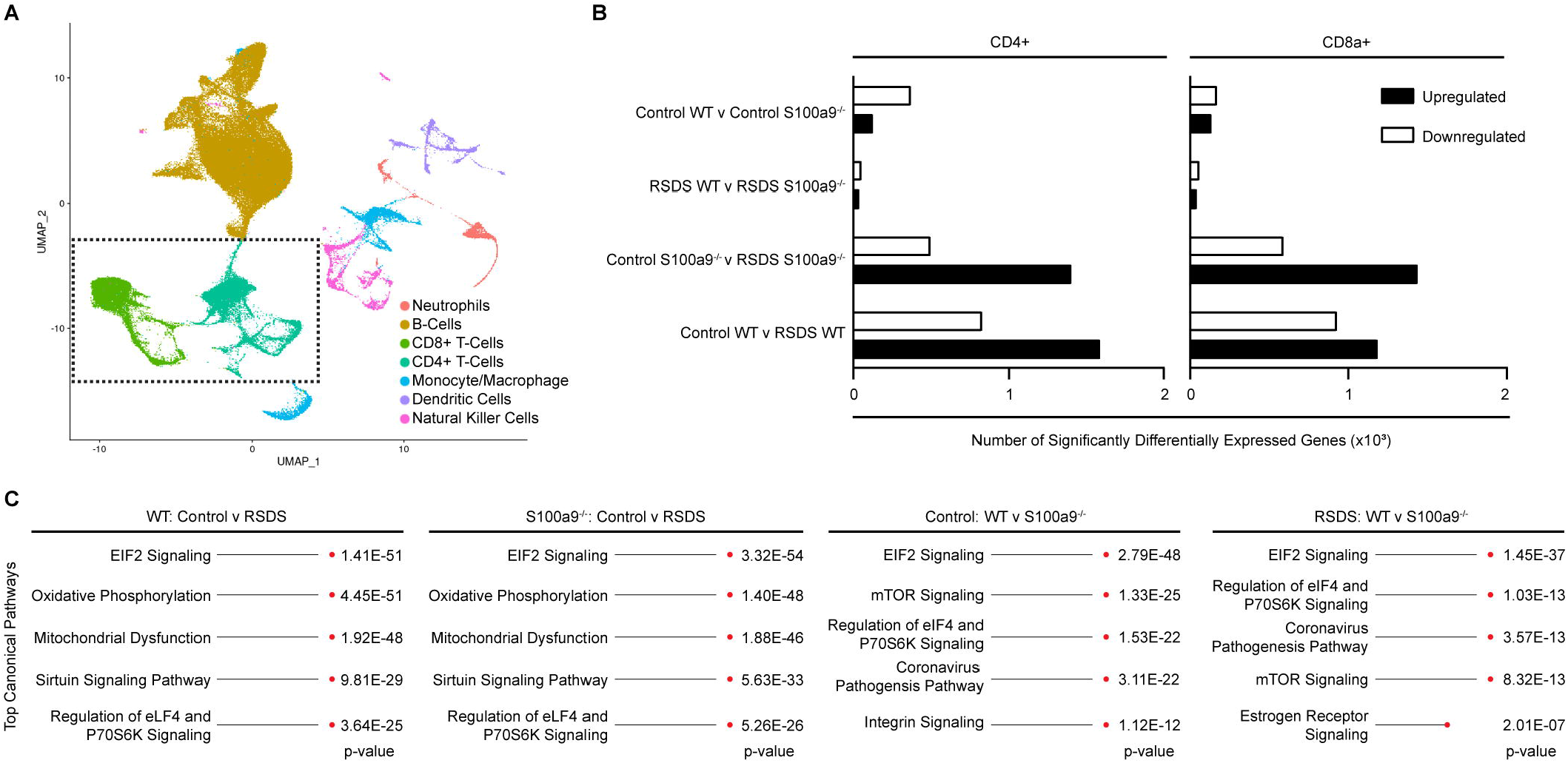
Loss of S100a9 perturbs T-lymphocyte gene expression similar to RSDS. WT and S100a9^-/-^ mice were run through RSDS, after which total splenocytes were isolated from these animals and assessed by single cell RNA sequencing. **A**. Seven primary cell population clusters were identified within the uniform manifold approximation and projection (UMAP). **B**. Quantification of number of significant differentially regulated genes among comparisons of genotype and psychological trauma. **C**. Ingenuity pathway analysis of differentially regulated genes among comparisons of genotype and psychological trauma in CD8^+^ and CD4^+^ T-lymphocytes (not shown because identical to CD8^+^).

Understanding that the mitochondrial redox environment is tightly coupled to metabolism (43), we next examined metabolism in the context of RSDS. RSDS significantly enhanced the basal respiration, spare respiratory capacity, and maximum respiration in pan T-lymphocytes from WT animals (**Figure 3B-E**), suggesting enhanced mitochondrial metabolism in these cells after psychological trauma. Interestingly, the loss of S100a9 appeared to increase these same parameters in T-lymphocytes at baseline, with RSDS having little to no effect on these cells regarding these metabolic parameters (**Figure 3B-E**). Examining the metabolic state further, we found that loss of S100a9 had a significant enhancement of both glycolytic and mitochondrial metabolism in purified CD8^+^ T-lymphocytes (**Figure 3F-O**) with only modest effects in purified CD4^+^ T-lymphocytes (**Supplemental Figure 5A-J**). These metabolic alterations with the loss of S100a9 cannot be explained by a difference in mitochondrial mass or proliferative capacity, as these were unchanged between WT and S100a9^-/-^ T-lymphocytes (**Supplemental Figure 5K-M**). Together, these data support that calprotectin plays a major metabolic homeostatic role in T-lymphocytes with loss of this protein leading to significant metabolic and redox shifts in these cells.

### Loss of S100a9 alone mimics RSDS-like gene expression patterns in T-lymphocytes

Due to S100a9 significantly impacting mitochondrial superoxide levels, inflammation, and metabolism in T-lymphocytes from RSDS mice, we performed single-cell RNA sequencing analysis on splenocytes from WT and S100a9^-/-^ control and RSDS mice (3 mice in each group; 12 mice total) to obtain a greater understanding of the gene expression changes in T-lymphocytes due to these perturbations. After compiling the data using the uniform manifold approximation and projection (UMAP) analysis (**Figure 4A**), we identified cell type clusters utilizing the *Tabula Muris* (40). Examining the differential gene expression of the CD4^+^ and CD8^+^ T-lymphocyte clusters, we found that RSDS induced a robust alteration in the genetic signature of both WT and S100a9^-/-^ T-lymphocytes (**Figure 4B-C**). Intriguingly, pathway analysis identified identical top canonical pathways altered in both WT and S100a9^-/-^ T-lymphocytes after RSDS (**Figure 4C**), with translation regulation and mitochondrial function being disrupted in both genotypes. Furthermore, no differences were noted between CD4^+^ and CD8^+^ T-lymphocytes, suggesting the changes associated with RSDS or S100a9^-/-^ loss have a universal effect on T-lymphocytes. In the absence of RSDS, the loss of S100a9 alone appeared to significantly impacts translation regulation and metabolic signaling (**Figure 4C**), thus suggesting that S100a9^-/-^ T-lymphocytes possess a phenotype at baseline similar to that of WT T-lymphocytes from an animal exposed to RSDS. This may explain why S100a8 and S100a9 were the top upregulated genes in our previous analysis of WT RSDS T-lymphocytes (19), as they may act as critical regulators of the processes disrupted by RSDS. Taken together, RSDS significantly impacts anabolic and metabolic genetic pathways in T-lymphocytes, while the loss of S100a9^-/-^ shifts T-lymphocytes into an RSDS-like state even in the absence of psychological trauma.

## Discussion

Calprotectin has been reported to primarily act as a pro-inflammatory protein, and its involvement in inflammatory diseases such as cancer, rheumatoid arthritis, psoriasis, endo-toxin induced shock, and obesity are well defined (20, 21, 23, 44-51). Furthermore, calprotectin is being investigated as a potential clinical biomarker for several inflammatory bowel, periodontal, autoimmune, and infectious diseases (including COVID-19) because of how tightly correlated the protein is with inflammatory profiles in these disease states (52-55). Additional evidence of the pro-inflammatory role of calprotectin is supported by the pharmacological inhibition of the protein using paquinimod, which demonstrates anti-inflammatory and beneficial effects in several of the aforementioned inflammatory pathologies (33, 56-60). However, examinations of calprotectin in the context of mental health are scarce.

To our knowledge, only three reports exist that have identified S100a8 and S100a9 upregulation in models of psychological stress, and all three reported the upregulation only occurring in regions of the brain (41, 61, 62). Of these three, only one attempted to examine the mechanistic role of calprotectin (62). In that work, Gong *et al*. examined the phenotypic effects of centrally administered recombinant calprotectin or paquinimod in a mouse model of chronic unpredictable stress (CUS). In line with our original hypothesis, they observed centrally infused calprotectin exacerbated depressive-like behavior and neuroinflammation, while pharmacological inhibition of calprotectin was shown to be beneficial in their animal model. Moreover, paquinimod was shown to directly attenuate reactive oxygen species production from a cultured microglial cell line, which further supported an anti-inflammatory effect of calprotectin inhibition (62). This singular study supports the notion that central calprotectin plays a pro-inflammatory and pro-psychopathological role after psychological stress, however, our findings suggest the opposite.

The disparity in these results may be due to several factors. First, we utilized different models of stress induction, which have different timelines, stressors, and phenotypes. The differences in our findings may demonstrate the complexity of calprotectin expression among psychopathologies, and a“one size fits all” approach may not be accurate. Second, our primary focus was on the effects of calprotectin in peripheral T-lymphocyte inflammation, while the others examined the consequences in the brain. It is quite possible calprotectin plays differential roles that are cell type dependent. While this may complicate therapeutic approaches with calprotectin, it does not minimize the possible role this protein may play as a biomarker of psychopathologies and their progression. Last, the dosage and route of administration of paquinimod differ between the studies. We utilized systemic infusion to block calprotectin, while the previous report centrally administered agents by intracerebroventricular injection. The dose we chose was also in the lower range reported in the literature to minimize off-target effects (33). Therefore, while the findings between these studies may seem contrasting, they may in fact highlight important nuances of calprotectin expression and function during psychological stress. Calprotectin is commonly referred to as a proinflammatory protein and a damage associated molecular patten due to its reported ability to activate toll-like receptor 4 (20). While our work presented herein challenges this canonical proinflammatory characteristic of calprotectin, others have also found that S100a8 and S100a9 provide protective functions in various contexts. For example, S100a8 administration was shown to induce anti-inflammatory IL-10 and protect against acute lung injury (63). In another study, loss of S100a9 potentiated the development of autoimmunity in a mouse model of lupus, suggesting S100a9 played a protective role in this context (64). An additional report found that S100a9 deficient neutrophils, macrophages, and dendritic cells all demonstrated differential cytokine expression in a model of atherosclerosis (65). Because not all cell types showed a similar pattern of inflammation, the authors (as well as others) concluded that the post-translational, cellular, and microenvironment contexts must play a significant role in how S100a8 and S100a9 function regarding inflammation (65, 66). Given that our work presented herein is the first examination of calprotectin in T-lymphocytes, it appears that under the context of psychological trauma that these proteins play a protective and anti-inflammatory role.

While we previously demonstrated calprotectin gene expression in T-lymphocytes (19), the work presented herein reports the first confirmed functional roles for the protein in these adaptive immune cells. First, it appears that S100a9 (and S100a8 since this protein is also lost with the knock-out of S100a9) play a significant role in the maintenance of metabolic homeostasis. As we observed in the absence of psychological trauma, both mitochondrial and glycolytic metabolism are greatly enhanced in T-lymphocytes isolated from S100a9^-/-^ mice. How calprotectin regulates T-lymphocyte metabolism remains unknown, but we hypothesize the mechanism may involve intracellular calcium sequestration. It is well established that calcium signaling plays an essential role in T-lymphocyte activation, proliferation, differentiation, and metabolism (67-69). Additionally, mitochondrial calcium uptake enhances mitochondrial energy output (70-73), which has been shown to be essential for T-lymphocyte polarization shifts from naive, activated, and memory states (43, 74-76). With this, calcium must be tightly regulated and controlled to avoid aberrant activation or differentiation, and calprotectin may act as an intracellular calcium sequestration protein to serve this purpose.

Calprotectin is able to bind 6 calcium ions (77), so it may act as a buffer to inhibit excess intracellular calcium signaling during T-lymphocyte activation. This hypothesis is supported by our data demonstrating that loss of calprotectin produced a hyper-activated T-lymphocyte state with pronounced metabolism, inflammation, and redox consequences.

Our original hypothesis that calprotectin may be perpetuating the negative inflammatory and redox consequences of RSDS in T-lymphocytes stemmed from our original observation that T-lymphocyte expressed calprotectin highly correlated with mitochondrial superoxide levels after RSDS (19). Given the previously discussed pro-inflammatory descriptions of calprotectin, it was logical to assume this protein was playing a similar role in the RSDS context. However, our results show quite the opposite, and suggest calprotectin may be playing a protective role in these cells. This suggests that calprotectin expression may be in response to the altered redox or inflammatory environments, as opposed to the cause. This concept is supported by early work in our laboratory examining the consequences of enhanced mitochondrial superoxide in T-lymphocytes. Reviewing this previous work where we used a mouse model of T-lymphocyte specific manganese superoxide dismutase knockout (to create an animal with uncontrolled mitochondrial superoxide in T-lymphocytes), we found that S100a8 and S100a9 were indeed elevated as assessed by an Affymetrix gene array (78). These data suggest that calprotectin upregulation in T-lymphocytes is downstream to mitochondrial redox changes, and may serve as a protective checkpoint necessary to prevent excessive activation or inflammation from T-lymphocytes. Future studies will examine if normalization of the mitochondrial redox environment in T-lymphocytes following RSDS is sufficient and/or necessary to restore normal calprotectin expression and inflammation in these cells.

Our data demonstrate a strong mitochondrial reaction in T-lymphocytes after RSDS. In addition to our metabolic and redox data, this was most supported by our single cell RNA sequencing data showing the top canonical gene pathways disrupted in these cells were related to mitochondrial function. The vast majority of these involved genes (94 in total) encode for the five major complexes of the electron transport chain. Intriguingly, of these 94 genes, all nuclear encoded mitochondrial genes were up-regulated, while all mitochondrial encoded mitochondrial genes were downregulated. This dichotomy suggests potential dysfunction within the mitochondria given their decreased abundance of transcripts, with a compensatory upregulation of nuclear encoded transcripts to counterbalance this defect. This exact same phenomenon is observed with the loss of S100a9 in T-lymphocytes even in the absence of psychological trauma, which demonstrates the previously undescribed importance of this protein in the maintenance of T-lymphocyte mitochondrial homeostasis. These similar genetic patterns are likely not random, but explain why S100a8 and S100a9 are two of the most upregulated genes in T-lymphocytes after RSDS. Our data suggest these proteins regulate the T-lymphocyte processes altered by psychological trauma, thus the loss of calprotectin alone leads to the net effect of increased cellular metabolism, enhanced mitochondrial superoxide production, and elevated inflammation similar to that of RSDS.

While these data provide new insight into psychological trauma-induced inflammation, this study is not without limitations. First, the mice utilized for these studies are constitutive S100a9 knock-outs, which limits our ability to make cell-type specific conclusions regarding systemic processes (*e*.*g*., behavior or circulating inflammation). At the time of this work, no conditional S100a9 knock-out mouse had been developed. This does not diminish the findings of this study, but encourages the development of a conditional S100a9 knock-out animal for more nuanced studies. Second, we have not performed a calprotectin rescue experiment in S100a9^-/-^ mice. We did attempt these studies using osmotic minipumps similar to that of paquinimod, but were unable to verify calprotectin in circulation of S100a9^-/-^ mice. At this time, we are unsure if this is due to a rapid breakdown of this protein when infused or an unknown technical limitation. Furthermore, the infusion of extracellular calprotectin will not restore intracellular protein in T-lymphocytes. This is additional incentive to develop a conditional S100a9^-/-^ mouse model where we can truly address the role of intracellular versus extracellular calprotectin on T-lymphocytes. Last, most studies herein were performed on male mice. This is due the inability of females to be incorporated into the standardized RSDS paradigm. We have successfully incorporated females using an adapted version with juvenile experimental mice, but utilizing mice at this young age poses numerous other challenges. We are currently adapting other established methods of female RSDS into our laboratory (79, 80), and hope to follow up these studies with S100a9^-/-^ female mice.

In summary, we put forth data supporting a protective role for calprotectin in T-lymphocytes after psychological trauma. While “protection” is in the protein’s namesake, this term was likely given to calprotectin due to its canonical ability to sequester calcium and metals, which protects from bacterial and other pathogen infections. These established roles for calprotectin primarily come from studies with neutrophils, where calprotectin makes up approximately 60% of neutrophil cytoplasmic protein, is rapidly excreted during times of infection, and is expressed at over 3 orders of magnitude the levels expressed in T-lymphocytes (which we suppose may be the reason it has not been described in T-lymphocytes to date). Our findings pave a new road for this protein in the context of T-lymphocytes and other cells where the role for calprotectin remains undefined.

## Acknowledgements

This work was supported by the National Institutes of Health (NIH) R01HL158521 (AJC), F30HL154535 (SKE), and R01MH118239 (VIV). The University of Nebraska Medical Center Genomics Core Facility receives partial support from the National Institute for General Medical Science (NIGMS) INBRE - P20GM103427-19, as well as the National Cancer Institute The Fred & Pamela Buffett Cancer Center Support Grant-P30CA036727. This publication’s contents are the sole responsibility of the authors and do not necessarily represent the official views of the NIH or NIGMS.

## Author Contributions

CMM and AJC designed research studies. All authors conducted experiments, acquired data, and/or performed analyses. CM and AJC wrote the manuscript. AJC provided experimental oversight.

## Conflict of Interest

The authors declare no competing financial interests in relation to the work described.

## Notes

**Conflict of Interest Statement:** The authors have declared that no conflict of interest exists.

### Competing Interest Statement

The authors have declared no competing interest.

