## Supplemental Data for "S100a9 Attenuates Inflammation during Repeated Social Defeat Stress"

**Affiliations:**
[1]Department of Cellular and Integrative Physiology, University of Nebraska Medical Center, Omaha, NE, United States
[2]Department of Pathology and Microbiology, University of Nebraska Medical Center, Omaha, NE, United States
[3]Department of Psychiatry and Behavioral Sciences, Texas A&M University, Bryan, TX, United States
[4]Department of Medical Physiology, Texas A&M University, Bryan, TX, United States
[5]Department of Psychiatry, University of Arizona, Phoenix, AZ, United States
[6]Lieber Institute for Brain Development, Johns Hopkins University, Baltimore, MD, United States

***Corresponding author:** Adam J. Case, PhD
Associate Professor
Department of Psychiatry and Behavioral Sciences
Department of Medical Physiology
8447 Riverside Pkwy
MREB2 3414
Bryan, TX 77807


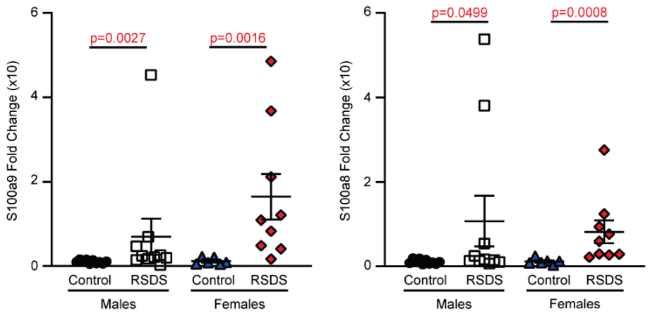


**Supplemental Figure 1. Elevated S100a8 and S100a9 gene expression are observed in T-lymphocytes from juvenile RSDS male and female mice.** Juvenile (3 week old) mice were run through RSDS, and after 1 month of recovery splenic T-lymphocytes were isolated. S100a9 and S100a8 mRNA levels assessed by real-time quantitative PCR in freshly isolated pan T-lymphocytes (n= 10 male controls, 10 male RSDS, 6 female control, 9 female RSDS). Significance by two-way ANOVA with Tukey's post hoc analysis throughout.

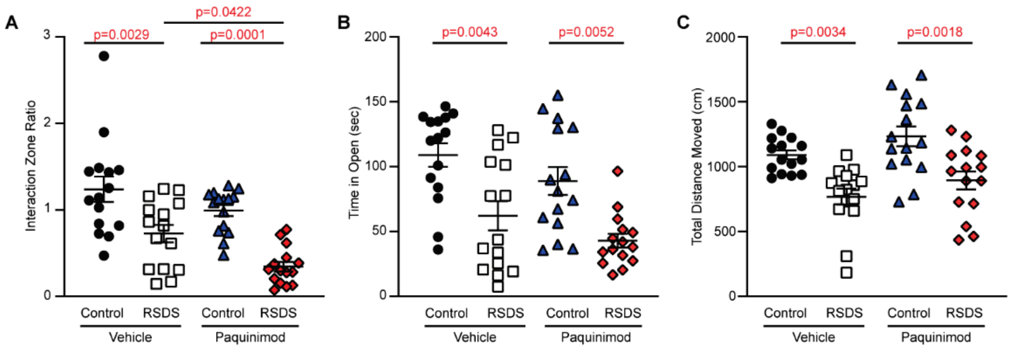


**Supplemental Figure 2. Pharmaceutical inhibition of calprotectin significant affects social behavior after RSDS.** WT mice were implanted with osmotic minipumps filled with either saline or paquinimod at a flow rate of 1.5 mg/kg/day three days prior to starting the RSDS protocol followed by behavior testing. **A**. Quantification of the interaction ratio from social interaction testing. **B-C.** Quantification of the time spent in open arms and total distanced moved from elevated zero maze testing (A-C: n = 15 saline control, 15 saline RSDS, 15 paquinimod control, 15 paquinimod RSDS). Statistics by two-way ANOVA with Tukey’s post hoc analysis throughout.

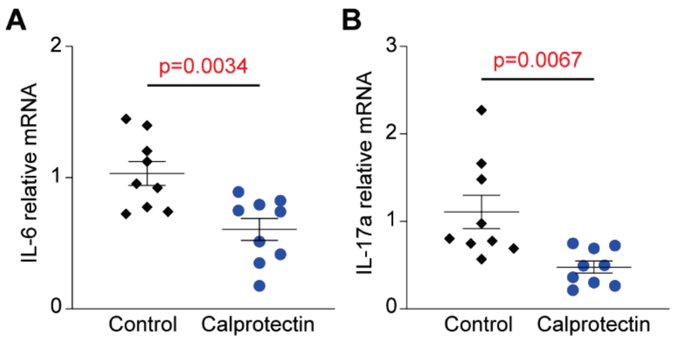


**Supplemental Figure 3. Calprotectin supplementation decreases T-lymphocyte pro-inflammatory cytokine expression.** Splenic T-lymphocytes were isolated from unstressed mice and cultured for 72 hours in the presence of calprotectin (1 µg/mL). **A-B**. IL-6 and IL-17A mRNA levels assessed by real-time quantitative PCR (n = 9 control, 9 calprotectin). Statistics by Student's t-test throughout.

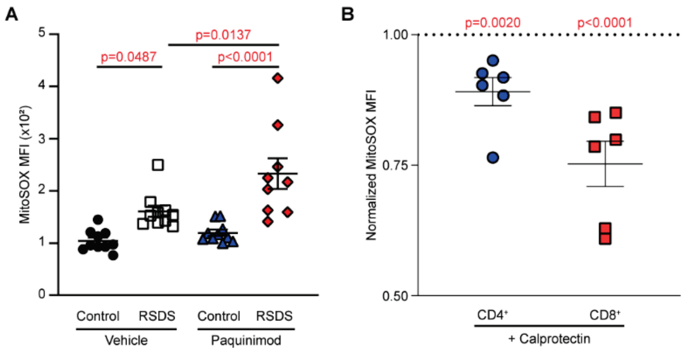


**Supplemental Figure 4. T-lymphocyte mitochondrial redox environment is significantly perturbed by calprotectin.** WT mice were implanted with osmotic minipumps filled with either saline or paquinimod at a flow rate of 1.5 mg/kg/day three days prior to starting the RSDS protocol followed by splenic T-lymphocyte isolation. **A**. MitoSOX Red mean fluorescent intensity (MFI) assessed by flow cytometry. Data normalized WT control T-lymphocytes to control for interexperimental variance (n= 10 WT controls, 10 WT RSDS, 10 S100a9$^{-/-}$ controls, 9 S100a9$^{-/-}$ RSDS). **B**. MitoSOX Red mean fluorescent intensity (MFI) assessed by flow cytometry in splenic T-lymphocytes isolated from unstressed mice and cultured for 72 hours in the presence of calprotectin (1 µg/mL). Data normalized to untreated CD4$^{+}$ or CD8$^{+}$ T-lymphocytes respectively (n= 6 CD4$^{+}$, 6 CD8$^{+}$). Statistics by paired Student's t-test or two-way ANOVA with Tukey's post hoc analysis where appropriate.

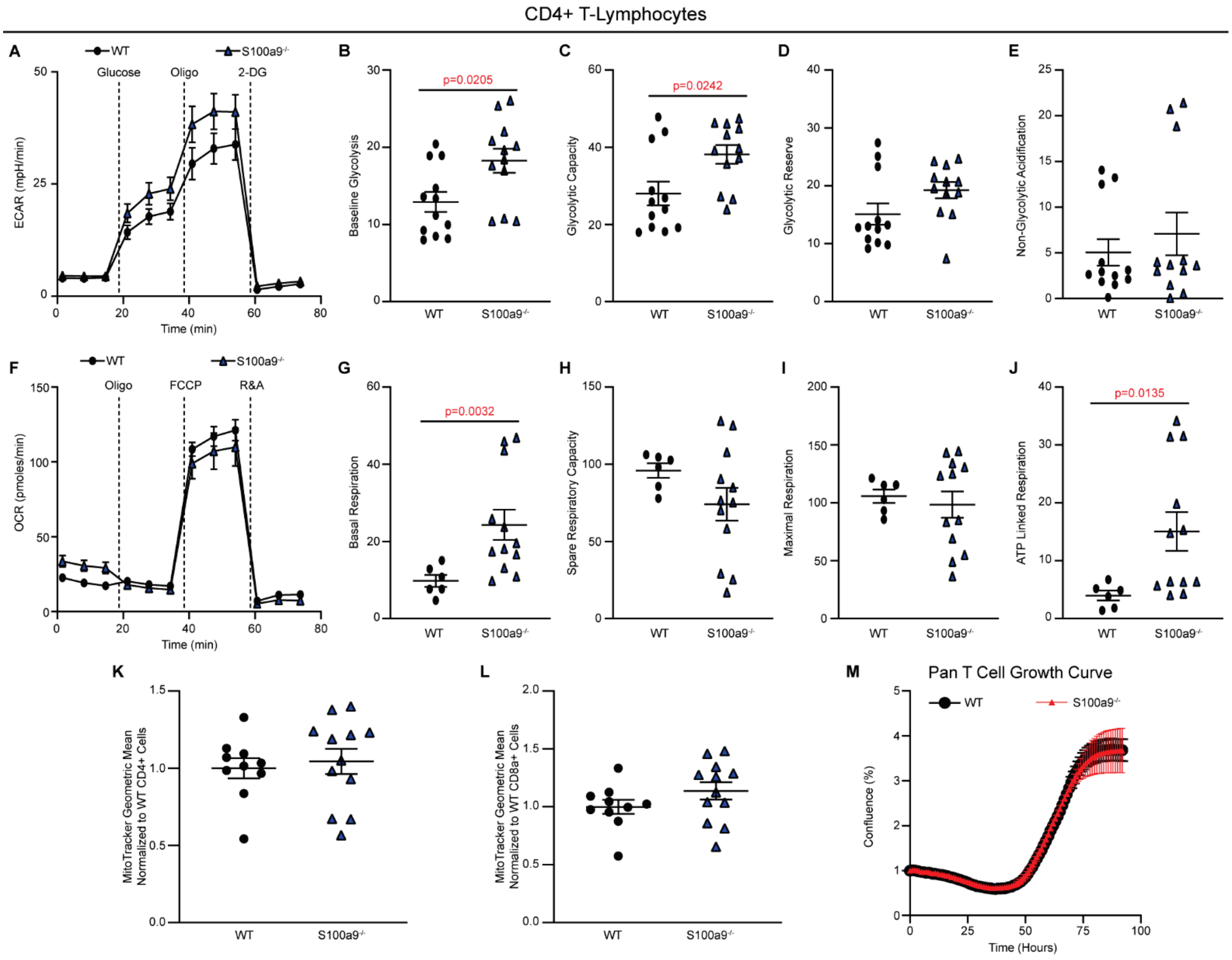


**Supplemental Figure 5. CD4⁺ T-lymphocytes are modestly affected by the loss of S100a9.** WT and S100a9$^{-/-}$ mice were run through RSDS, after which splenic T-lymphocytes were isolated from these animals. **A**. Representative extracellular acidification rate (ECAR) curve of a glycolysis stress test (n= 12 WT, 12 S100a9$^{-/-}$). **B-E**. Quantification of metabolic states from the glycolytic stress test. **F**. Representative oxygen consumption rate (OCR) curve of mitochondrial stress test (n= 6 WT, 12 S100a9$^{-/-}$). **G-J**. Quantification of metabolic states from the mitochondrial stress test. **K-L**. MitoTracker green fluorescence assessed by flow cytometry in CD4$^{+}$ and CD8$^{+}$ T-lymphocytes, respectively. (n= 10 WT, 12 S100a9$^{-/-}$). **M**. Growth curve of activated splenic T-lymphocytes assessed by Incucyte S3 live cell imaging. (n= 3 WT, 3 S100a9$^{-/-}$). Statistics by Mann-Whitney U-test throughout.

| Name | Sequence |
|---|---|
| 18s F | GCCCGAAGCGTTTACTTTGA |
| 18s R | TCATGGCCTCAGTTCCGAA |
| IL-2 F | TCTACAGCGGAAGCACAGC |
| IL-2 R | CCTGGGGAGTTTCAGGTTC |
| IL-6 F | GCTACCAAACTGGATATAATCAGGA |
| IL-6 R | CCAGGTAGCTATGGTACTCCAGAA |
| IL-17A F | CAGGGAGAGCTTCATCTGTGT |
| IL-17A R | GCTGAGCTTTGAGGGATGAT |
| TNFa F | CTGTAGCCCACGTCGTAGC |
| TNFa R | TTGAGATCCATGCCGTTG |

**Supplemental Table 1. SYBR Green oligonucleotide sequences used to measure cytokine gene expression change by quantitative real-time PCR.**
